# Dietary fatty acids promote sleep through a taste-independent mechanism

**DOI:** 10.1101/681635

**Authors:** Estelle Laure Sah Pamboro, Elizabeth B. Brown, Alex C. Keene

## Abstract

Consumption of foods that are high in fat contributes to obesity and metabolism-related disorders that are increasing in prevalence and present an enormous health burden throughout the world. Dietary lipids are comprised of triglycerides and fatty acids, and the highly palatable taste of dietary fatty acids promotes food consumption, activates reward centers in mammals, and underlies hedonic feeding. Despite a central role of dietary fats in the regulation of food intake and the etiology of metabolic diseases, little is known about how fat consumption regulates sleep. The fruit fly, *Drosophila melanogaster*, provides a powerful model system for the study of sleep and metabolic traits, and flies potently regulate sleep in accordance with food availability. To investigate the effects of dietary fats on sleep regulation, we have supplemented fatty acids into the diet of *Drosophila* and measured their effects on sleep and activity. We found that feeding flies a diet of hexanoic acid, a medium-chain fatty acid that is a by-product of yeast fermentation, promotes sleep by increasing the number of sleep episodes. This increase in sleep is dose-dependent and independent of the light-dark cues. Diets consisting of other fatty acids, including medium- and long-chain fatty acids, also increase sleep, suggesting many fatty acid types promote sleep. To assess whether dietary fatty acids regulate sleep through the taste system, we assessed sleep in flies with a mutation in the hexanoic acid receptor *Ionotropic receptor 56d*, which is required for fatty acid taste perception. We found that these flies also increase their sleep when fed a hexanoic acid diet, suggesting the sleep promoting effect of hexanoic acid is not dependent on sensory perception. Overall, these results define a role for fatty acids in sleep regulation, providing a foundation to investigate the molecular and neural basis for fatty acid-dependent modulation of sleep duration.

## Introduction

Sleep is a near universal behavior that is influenced by diverse environmental factors and life-history traits (Allada & Siegel, 2008; Joiner, 2016). While the function of sleep remains unclear, foraging strategies are a proposed factor underlying the robust inter-species variation in sleep duration throughout the animal kingdom (Capellini, Barton, McNamara, Preston, & Nunn, 2008; Keene & Duboue, 2018; Siegel, 2005). Further, diverse species acutely regulate sleep in accordance with nutrient availability, suggesting evolutionarily ancient interactions between sleep, foraging, and metabolic state (Keene & Duboue, 2018; M. Yurgel, Masek, DiAngelo, & Keene, 2014). Despite the prevalence of these interactions, the mechanistic basis for interactions between sleep and energy balance are largely unknown.

The fruit fly, *Drosophila melanogaster*, provides a powerful system to investigate the interrelationship between diet and sleep regulation (Chakravarti, Moscato, & Kayser, 2017; Griffith, 2013). Flies display all the behavioral hallmarks of sleep including prolonged immobility, elevated arousal threshold, and rebound following sleep deprivation (Hendricks et al., 2000; Shaw, Cirelli, Greenspan, & Tononi, 2000). Recent work has identified behavioral, electrophysiological, and metabolic states that associate with light and deeper sleep, suggesting distinguishable sleep stages are also present in fruit flies (Alphen, Yap, Kirszenblat, Kottler, & Swinderen, 2013; Faville, Kottler, Goodhill, Shaw, & van Swinderen, 2015; B. A. Stahl, Slocumb, Chaitin, DiAngelo, & Keene, 2017). In addition, genetic analyses suggest the molecular and neural circuit principles regulating sleep, as well as the functional effects of sleep loss, are highly conserved between flies and mammals (Donlea, 2017; Sehgal & Mignot, 2011). Therefore, flies present a highly tractable model for investigating how dietary factors modulate sleep.

Diet potently effects sleep, and the consumption of foods that are high in fat is thought to promote increased feeding and contributes to sleep-related disorders (Arble et al., 2015; Tan et al., 2015). Food is comprised of complex macronutrients including sugars, proteins, and fats. In flies, diet composition has been shown to impact sleep, but studies have primarily relied upon varying the concentrations of sugars and protein, or fully food-restricting animals (Catterson et al., 2010; Keene et al., 2010; Linford, Chan, & Pletcher, 2012; Linford, Ro, Chung, & Pletcher, 2015; Yamazaki et al., 2012). Flies have are capable of tasting a number of additional modalities including fatty acids, salt, amino acids, and carbonation (Fischler, Kong, Marella, & Scott, 2007; Masek & Keene, 2013; Toshima & Tanimura, 2012; Zhang, Ni, & Montell, 2013), but the effects of these factors on sleep regulation has not been explored.

Flies are attracted to fatty acids, which are sensed by a shared population of neurons that also detect sugar and have been implicated in sleep regulation (Masek & Keene, 2013). These neurons express the *Ionotropic Receptor 56d* (*IR56D*), which is required for fatty acid taste perception and presumably serves as a receptor for medium-chain fatty acids (Ahn, Chen, & Amrein, 2017; J. Tauber et al., 2017; Zappia et al., 2018). Fatty acids, as well as esters and alcohols, are a by-product of yeast fermentation and emitted from ripened fruit (Styger, Jacobson, & Bauer, 2011; Zhu et al., 2018). We have previously found that flies fed a diet of only medium-chain fatty acids live longer than starved flies (Masek & Keene, 2013), suggesting fatty acids provide nutrients, but their role in sleep regulation has not been tested. Here, we systematically characterize the effects diverse classes of fatty acids on fly sleep.

## Results

To determine the effects of dietary fatty acids on sleep, we quantified sleep duration and architecture in the *Drosophila* Activity Monitoring (DAM) system (Hendricks et al., 2000; Shaw et al., 2000). After an acclimation period, baseline sleep of mated female flies was measured over a 24 hr period, followed by 24 hrs of sleep measurements on a diet consisting of agar (starvation) or agar and hexanoic acid (Fig 1A). We first measured sleep on hexanoic acid (HxA), a 6-chain fatty acid that induces feeding behavior in *Drosophila* (Masek & Keene, 2013). Flies fed HxA slept significantly more during the daytime and nighttime than flies fed a diet of agar alone, revealing a sleep-promoting role for HxA (Fig 1B,C). The increase in sleep was a result of a greater number of sleep bouts, rather than an increase in bout length (Fig 1D,E). In addition to suppressing sleep, starvation results in hyperactivity (Keene et al., 2010; G. Lee & Park, 2004). We found that waking activity in flies fed a diet of HxA alone did not differ from agar-fed flies, suggesting that HxA does not ameliorate starvation-induced hyperactivity (Fig 1F). To determine the contributions of the light-dark cycle to the role of dietary HxA on sleep, we shifted the time of starvation to the onset of lights off and measured sleep for 24 hrs. In these flies, both nighttime and daytime sleep is greater than flies fed agar alone, fortifying the notion that dietary FA’s promote sleep (Fig 1G,H). Again, this longer sleep duration was a result of an increase in the number of sleep bouts, and not bout length (Fig 1I, Fig S1A). Therefore, dietary HxA selectively promotes sleep without affecting starvation-induced hyperactivity. To confirm the effects of HxA on survivorship, we measured starvation resistance in flies fed HxA or agar alone. In agreement with previous literature (Masek & Keene, 2013), we found that flies survive significantly longer upon the addition of HxA (Fig S2).

**Figure 1.**
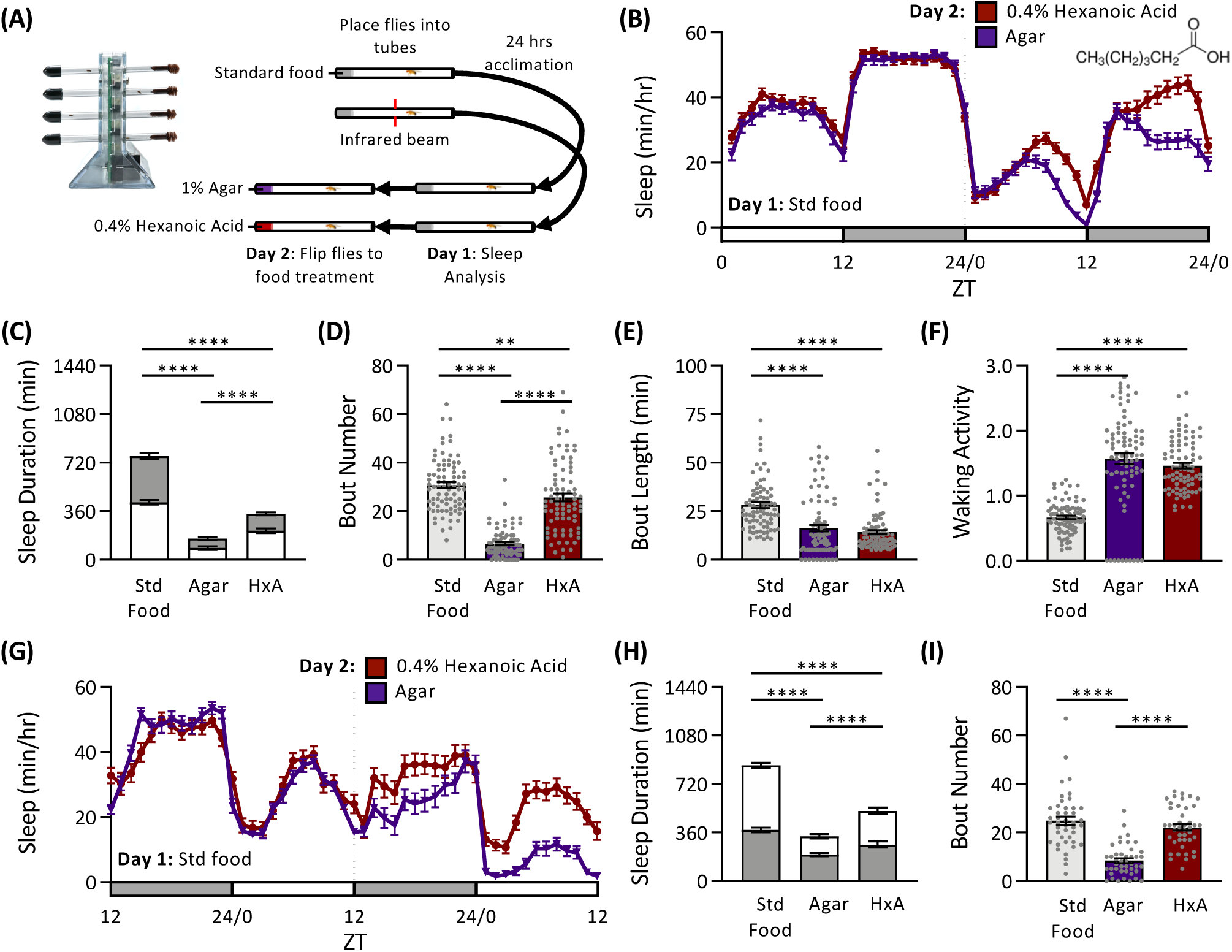
The addition of hexanoic acid to the diet partially rescues sleep loss. **(A)** Sleep traits were measured using the *Drosophila* Activity Monitoring (DAM) system. Individual flies were placed inside plastic tubes containing standard food at one end and a foam plug at the other. Flies were then allowed to acclimate for a minimum of 24hrs. At ZT0, baseline sleep and activity were measured on a standard food diet. At ZT0 of the following day, flies were transferred into tubes containing either agar or agar and 0.4% hexanoic acid (HxA), and again, sleep and activity were assessed. **(B)** Sleep profile of flies fed standard food on Day 1 and then either agar or HxA on Day 2. **(C)** Sleep duration is dependent on food treatment (two-way ANOVA: F_2,494_= 211.6, *P*<0.0001). Relative to agar, sleep significantly increases when HxA is added to the diet (*P*<0.0001). However, both treatments sleep significantly less than flies fed standard food (agar: *P*<0.0001; HxA: *P*<0.0001). **(D)** The number of sleep episodes is dependent on food treatment (one-way ANOVA: F_2,247_= 111.5, *P*<0.0001). Relative to agar alone, sleep bout number significantly increases when HxA is added to the diet (*P*<0.0001), but is significantly lower than when fed standard food (*P*=0.0091). **(E)** The length of each sleep episode is dependent on food treatment (one-way ANOVA: F_2,247_= 26.85, P<0.0001). Flies fed standard food have significantly longer bout lengths than flies fed a diet of either agar (*P*<0.0001) or HxA (*P*<0.0001), which are not significantly different from each other (*P*=0.5920). **(F)** Waking activity is dependent on food treatment (one-way ANOVA: F_2,247_= 78.5, *P*<0.0001). Flies fed standard food are significantly less active than flies fed a diet of either agar (*P*<0.0001) or HxA (*P*<0.0001), which are not significantly different from each other (*P*=0.3790). **(G-I)** In contrast to **(A)**, flies were transferred to their respective diet at ZT12. Sleep traits were measured either 24 hrs prior to or after this timepoint. **(G)** Sleep profile of flies fed standard food on Day 1 and then either agar or HxA on Day 2. **(H)** Sleep duration is dependent on food treatment (two-way ANOVA: F_2,312_= 96.57, *P*<0.0001). Relative to agar, sleep significantly increases when HxA is added to the diet (*P*<0.0001). However, both treatments sleep significantly less than flies fed standard food (agar: *P*<0.0001; HxA: *P*<0.0001). **(I)** The number of sleep episodes is dependent on food treatment (one-way ANOVA: F_2,126_=39.91, *P*<0.0001). Relative to agar alone, sleep bout number significantly increases when HxA is added to the diet (*P*<0.0001), but is not different from flies fed standard food (*P*=0.3417). For sleep profiles, white bars represent daytime (lighting is on), while gray bars represent nighttime (lighting is off). Hatched line indicates the time at which flies were transferred from standard food to their respective diet. Bar graphs display mean ± s.e.m. Gray circles represent measurements of individual flies. B-F: N = 83-84; G-I: N = 43. ** *P*<0.01; *** *P*<0.001; **** *P*<0.0001.

To investigate the effect of fatty acid concentration and sleep we compared sleep in flies fed agar alone or increasing concentrations of HxA. We found that sleep duration did not differ in in flies fed agar, 0.05%, or 0.1% HxA (Fig 2A). Flies fed a concentration of 0.4% and 0.7% HxA slept more than agar-alone flies but less than flies fed standard food, while feeding flies a diet consisting exclusively of 1.0% HxA resulted in sleep that was comparable in duration to flies fed standard food (Fig 2A). Similarly, we observed a concentration-dependent increase in the number of sleep episodes, with bout number significantly increasing beginning at 0.4% HxA (Fig 2B). Bout length was largely unaffected in flies fed HxA, except at a low concentration of 0.05% HxA, where it was increased in comparison to food- and agar-fed flies (Fig 2C). We also found that waking activity decreases with increasing concentrations of HxA, and at 1.0% HxA does not differ from flies fed standard food (Fig 2D). Therefore, the addition of HxA to the diet restores sleep to starved flies in a concentration-dependent fashion.

**Figure 2.**
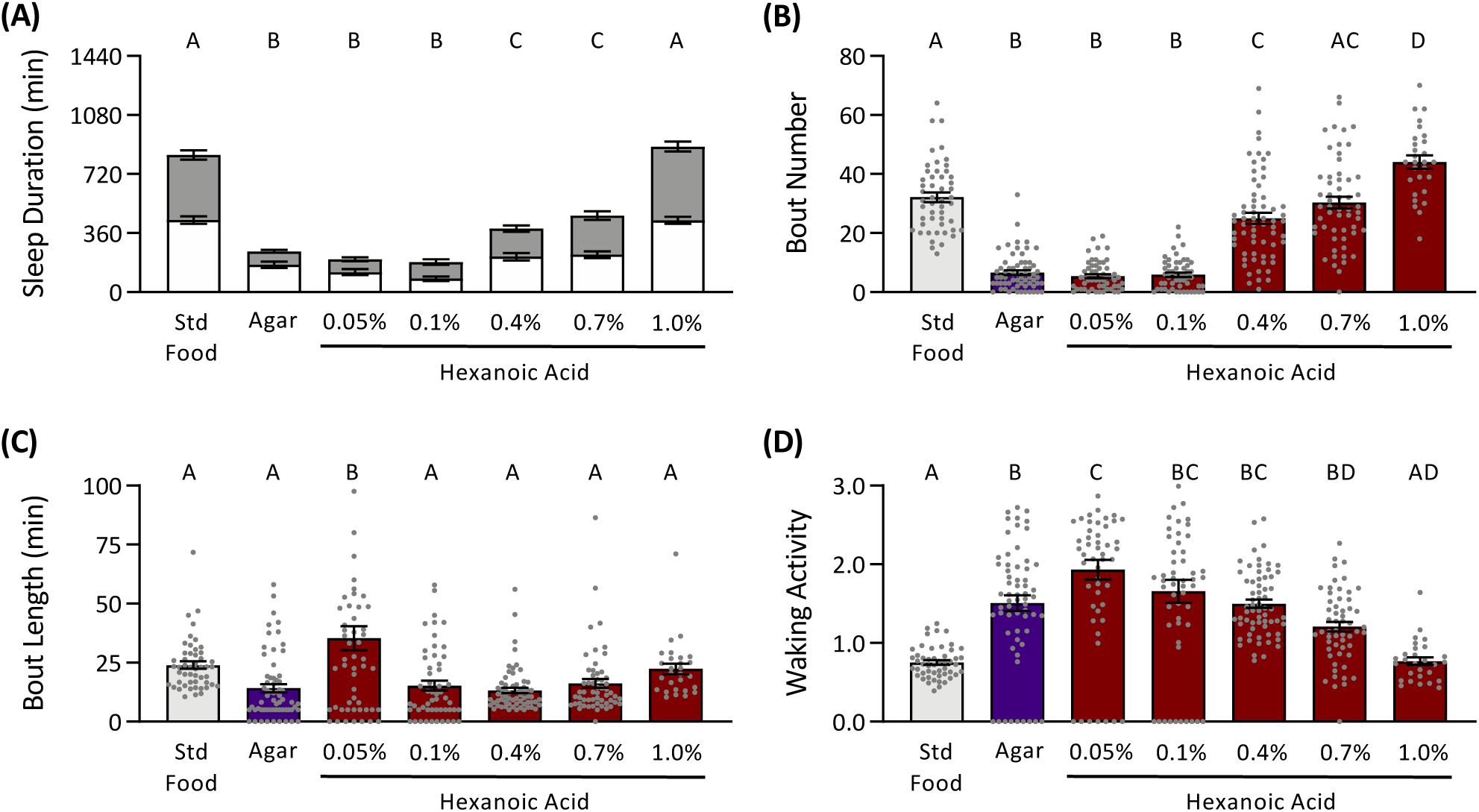
The modulation of sleep duration by hexanoic acid is dose-dependent. Hexanoic acid, ranging from a concentration of 0.05% to 1% was added to 1% agar and sleep traits were assessed. **(A)** There is a concentration-specific effect of hexanoic acid in the diet on sleep duration (two-way ANOVA: F_6,722_= 88.23, *P*<0.0001). Relative to agar, sleep duration significantly increases at concentrations at or above 0.4% HxA. Relative to standard food, sleep duration is significantly less, except at a concentration of 1% HxA, at which there is no difference. **(B)** There is a concentration-specific effect of hexanoic acid in the diet on the number of sleep episodes (one-way ANOVA: F_6,361_= 93.20, *P*<0.0001). Relative to agar, bout number significantly increases at concentrations at or above 0.4% HxA. Relative to standard food, bout number is significantly less, except at concentrations of 0.7% and 1% HxA, at which there is no difference. **(C)** There is a concentration-specific effect of hexanoic acid in the diet on the length of each sleep episode (one-way ANOVA: F_6,361_= 9.938, *P*<0.0001). Bout length significantly increases when flies are fed 0.05% HxA, relative to all other concentrations and food treatments. **(D)** There is a concentration-specific effect of hexanoic acid in the diet on waking activity (one-way ANOVA: F_6,361_= 19.41, *P*<0.0001). Relative to agar, there is no change in waking activity except at the lowest (0.05%) and highest (1%) concentrations. Relative to standard food, waking activity significantly increases, except at a concentration 1% HxA, at which there is no difference. Treatments that are significantly different from each other are represented with different letters *(P*<0.05). Bar graphs display mean ± s.e.m. Gray circles represent measurements of individual flies. N = 28-64.

To determine whether HxA was sufficient to promote sleep in otherwise fed flies, we assessed sleep when HxA was added to sucrose or to a standard *Drosophila* diet. Flies fed HxA and sucrose sleep longer than flies fed sucrose alone during the daytime and nighttime (Fig 3A,B). Similar to HxA alone, this increase in sleep was a result of to an increased number of sleep bouts rather than in increase in their length (Fig 3C,D). However, there was no change in waking activity (Fig 3E), suggesting that the increased sleep in HxA fed flies is not caused by lethargy. When HxA was added to standard food, there was no change in sleep duration, nor in any sleep traits (Fig 3 F-J), suggesting the sleep promoting role of HxA is not additive to a diet rich in sugar and yeast. Our findings that supplementation of HxA to sucrose, but not standard food, promotes sleep suggests that the addition of fatty acids to the diet selectively promotes sleep on nutrient-poor food and not on nutrient-rich food.

**Figure 3.**
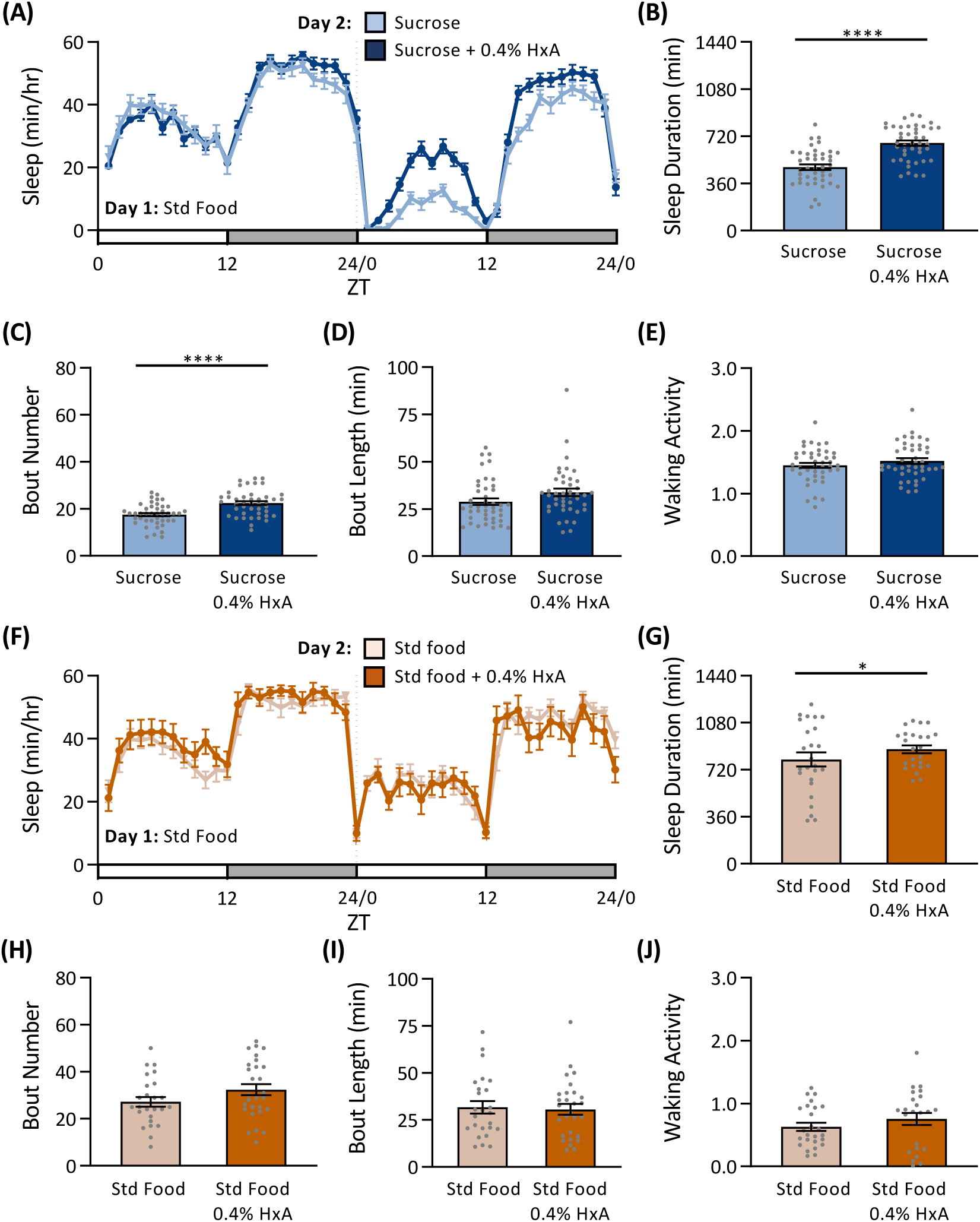
The sleep promoting effects of hexanoic acid is dependent on food substrate. Hexanoic acid was added to either 5% sucrose or standard food and sleep traits were assessed. **(A)** Sleep profile of flies fed standard food on Day 1 and then either maintained on 5% sucrose or 5% sucrose + 0.4% HxA on Day 2. **(B)** Sleep duration significantly increases when HxA is added to a sucrose-only diet (two-way ANOVA: F_1,168_= 38.86, *P*<0.0001). These differences persist during both the day (*P*=0.0001) and night (*P*<0.0001). **(C)** The number of sleep episodes significantly increases when HxA is added to a sucrose-only diet (t-test: t_84_= 4.481, *P*=0.0001). **(D)** There is no change in the length of each episode (t-test: t_84_= 1.846, *P*=0.0684). **(E)** There is also no change in waking activity (t-test: t_84_= 1.161, *P*=0.2491). **(F)** Sleep profile of flies fed standard food on Day 1 and then either maintained on standard food or standard food + 0.4% HxA on Day 2. **(G)** Although there is a significant effect of adding HxA to a standard food diet (two-way ANOVA: F_1,94_=5.664, *P*=0.0193), post hoc analyses revealed no difference either during the day (*P*=0.3438) or the night (*P*=0.0853). **(H)** There is no change in the number of sleep episodes when HxA is added to a sucrose-only diet (t-test: t_51_= 1.629, *P*=0.1096). (**I)** There is no change in the length of each episode (t-test: t_51_= 0.255, *P*=0.7998). (**J)** There is also no change in waking activity (t-test: t_51_= 1.076, *P*=0.2875). For sleep profiles, white bars represent daytime (lighting is on), while gray bars represent nighttime (lighting is off). Hatched line indicates the time at which flies were transferred from standard food to their respective diet. Bar graphs display mean ± s.e.m. Gray circles represent measurements of individual flies. A-E: N = 25-28; F-J: N = 43. ** *P*<0.01; *** *P*<0.001; **** *P*<0.0001.

It is possible that fatty acids generally promote sleep, or that it is selective to HxA. To differentiate between these possibilities, we tested sleep in flies fed individual FAs ranging from 3 carbons (3C; propionic acid) to 10 carbons (10C; decanoic acid). Flies fed fatty acids with a length of 6 carbons or greater increased sleep, with the most robust effect in flies fed 7-10C fatty acids (Figure 4A). Interestingly, while 6-10C fatty acids all promoted sleep relative to starved flies, they had different effects on sleep architecture. Fatty acids of 3-5C significantly increased bout number to a moderate degree, although this increase did not result in an overall increase in sleep duration (Figure 4B). Further, compared to starved flies, feeding flies 6-9C fatty acids also significantly increased bout number, but decanoic acid (10C) did not (Figure 4B). With regard to the length of each sleep bout, we observed a significant increase in bout length only in flies fed long-chain (9C and 10C) fatty acids (Figure 4C). Relative to both standard food and agar, bout length was unchanged in flies fed either short- or long-chain fatty acids. Further, all fatty acids tested generally produced a similar degree of hyperactivity as that of agar-fed flies (Figure 4D), with the exception of heptanoic acid (7C). The addition of this fatty acid resulted in waking activity that was intermediate between standard food and agar. The diverse effects of fatty acids on sleep architecture raise the possibility that different mechanisms underlie the sleep-promoting effects of fatty acids on sleep.

**Figure 4.**
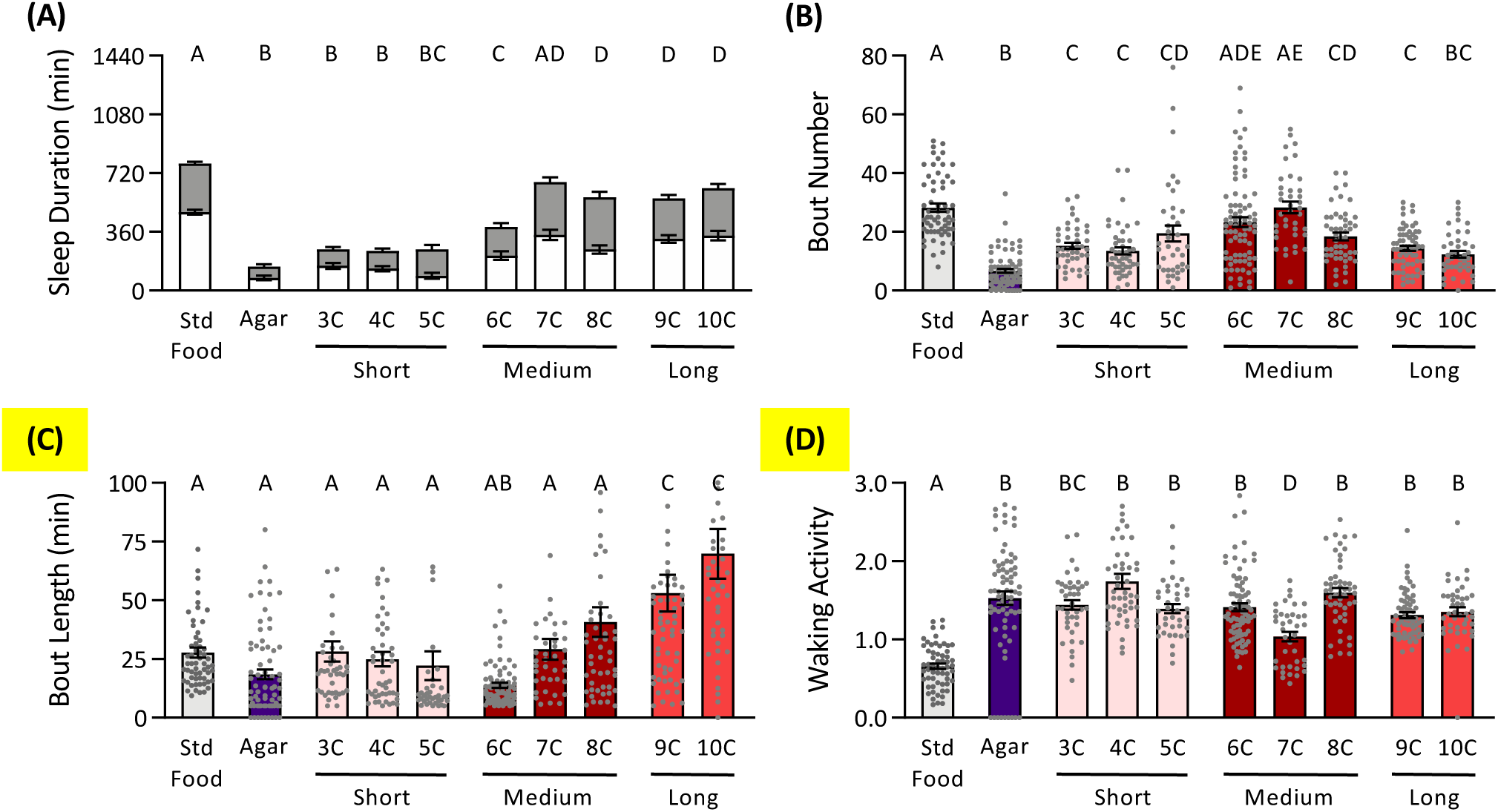
Medium- and long-chain dietary fatty acids promote sleep. Saturated fatty acids ranging from 3 carbon chains to 10 carbon chains were added to 1% agar at a concentration of 0.4%. **(A)** There is a chain length-specific effect of fatty acids on sleep duration (two-way ANOVA: F_9,1008_= 55.70, *P*<0.0001). Relative to agar, sleep duration significantly increases when medium or long-chain fatty acids are added to the diet. **(B)** There is a chain length-specific effect of fatty acids on the number of sleep episodes (one-way ANOVA: F_9,525_= 25.70, *P*<0.0001). Relative to agar, bout number significantly increases regardless of carbon chain length. **(C)** There is chain length-specific effect of fatty acids on the length of each sleep episode (one-way ANOVA: F_9,525_= 11.42, *P*<0.0001). There is a significantly increase in bout length only when long-chain fatty acids are added to the diet. **(D)** There is a chain length-specific effect of fatty acids on waking activity (one-way ANOVA: F_9,525_= 22.98, *P*<0.0001). Relative to standard food, waking activity significantly increases regardless of carbon chain length. However, this increase in waking activity is not significantly different from that of starved flies. Treatments that are significantly different from each other are represented with different *letters (P*<0.05). Bar graphs display mean ± s.e.m. Gray circles represent measurements of individual flies. N = 38-80.

Hexanoic acid and sugars are detected by shared peripheral sensory neurons that known to be sleep promoting (Ahn, Chen, & Amrein, 2017; J. M. Tauber et al., 2017). Therefore, it is possible that fatty acids promote sleep by providing nutrients, or through the activation of these sensory neurons. To differentiate between these possibilities, we measured sleep in flies lacking the HxA receptor, *Ionotropic Receptor 56d* (*IR56D*). Overall sleep duration, architecture, and waking activities did not differ between wild type and flies lacking *IR56D* indicating that *IR56D* is dispensable for the regulation of sleep and activity. We found that supplementing agar with HxA increased sleep in both control and *IR56D* null flies, suggesting that taste perception of HxA is not required for its sleep promoting effects (Fig 5A). Similarly, sleep architecture did not differ between control and *IR56D* null flies, as both control and *IR56D* null flies significantly increased the number of sleep bouts without increasing bout length (Fig 5B,C). Further, the addition of HxA to the diet did not change waking activity in either the control nor the *IR56D* null flies (Fig 5D). To determine whether *Ir56d* neurons are involved in sleep regulation, we blocked neurotransmitter release from *IR56D*-expressing neurons using tetanus toxin (Sweeney, Broadie, Keane, Niemann, & Kane, 1995) and observed similar results (Fig 5E-H): that silencing *IR56D*-expressing neurons has no effect on the regulation of sleep and activity. Therefore, peripheral taste perception of HxA is dispensable for the sleep promoting effects of dietary HxA.

**Figure 5.**
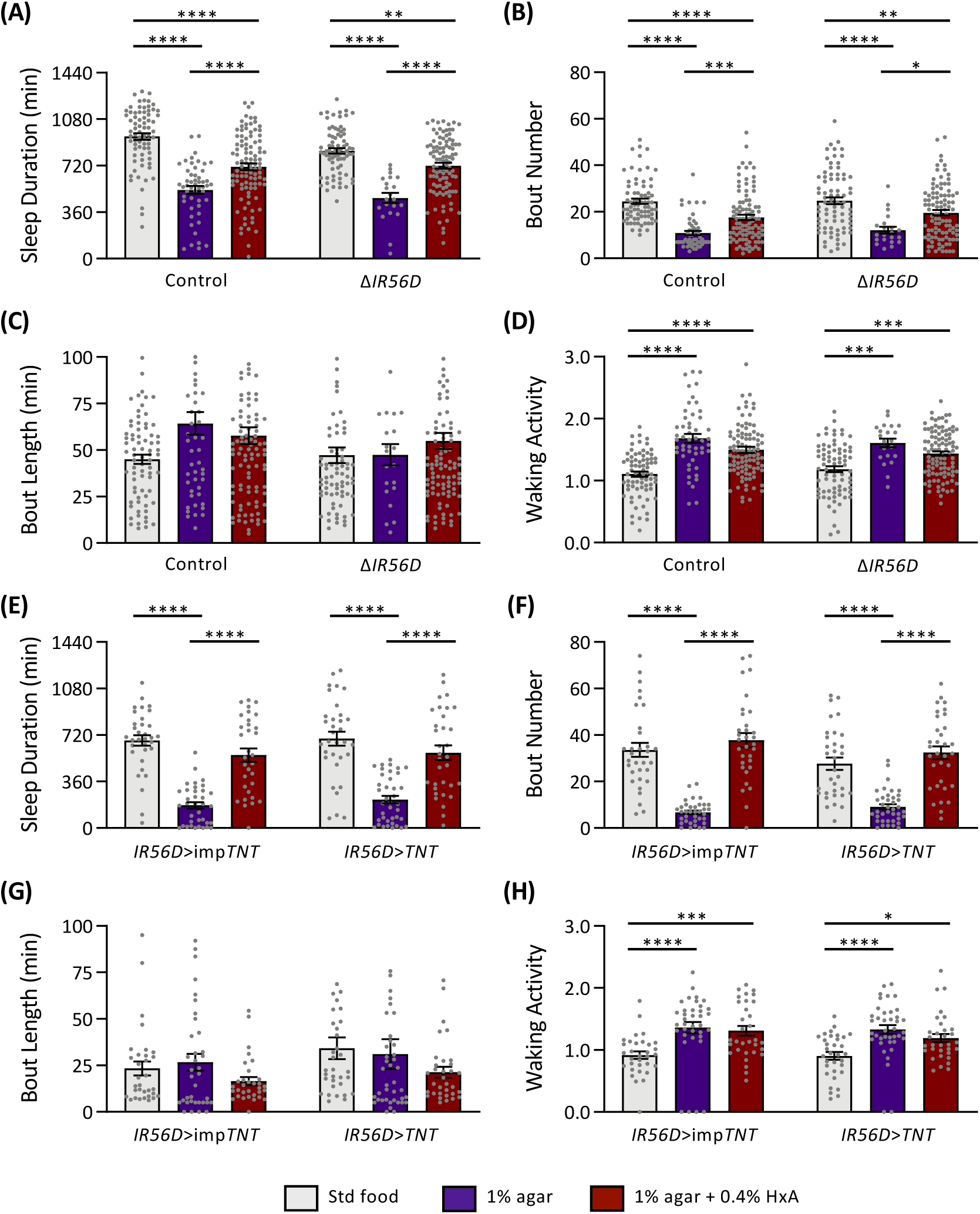
The gene *IR56D* is not required for fatty acid-dependent sleep regulation. Sleep traits were assessed in flies either lacking the IR56D gene or in flies whose neurons expressing the *IR56D* gene were silenced. **(A)** There is a significant effect of genotype on sleep duration in *IR56D* mutant flies (two-way ANOVA: F_1,418_= 4.954, *P*=0.0266), but post hoc analyses revealed no difference for each food type (Std food: *P*=0.2774; Agar: *P*=0.7933 HxA: *P*=0.3056). There is also an effect of food treatment (two-way ANOVA: F_2,418_= 70.96, *P*<0.0001). Relative to agar, both control and *IR56D* mutants significantly increase sleep duration when HxA is added to the diet (w^1118^: *P*<0.0001; Δ*IR56D*: *P*<0.0001), and both genotypes sleep significantly less than flies fed standard food (w^1118^: *P*<0.0001; Δ*IR56D*: *P*=0.0019). **(B)** There is no effect of genotype on the number of sleep episodes in *IR56D* mutant flies (two-way ANOVA: F_1,418_= 0.9479, *P*=0.3308), but the effect of food treatment remains (two-way ANOVA: F_2,418_= 22.72, *P*<0.0001). Relative to agar, both control and *IR56D* mutants significantly increase sleep bout number when HxA is added to the diet (w^1118^: *P*=0.0008; Δ*IR56D*: *P*=0.0104), however it is significantly less than flies fed standard food (w^1118^: *P*<0.0001; Δ*IR56D*: *P*=0.0104). **(C)** There is no effect of genotype on the length of each sleep episode in *IR56D* mutant flies (two-way ANOVA: F_1,418_= 1.763, *P*=0.1850), nor is there an effect of food treatment (two-way ANOVA: F_2,418_= 3.187, *P*=0.0623). **(D)** There is no effect of genotype on waking activity (two-way ANOVA: F_1,418_= 0.1469, *P*=0.7017), but the effect of food treatment remains (two-way ANOVA: F_2,418_= 39.38, *P*<0.0001). Relative to agar, there is no change in waking activity in either control or *IR56D* mutants upon addition of HxA (w^1118^: *P*=0.2770; Δ*IR56D*: *P*=0.2091), but is significantly higher than on standard food (w^1118^: *P*<0.0001; Δ*IR56D*: *P*=0.0003). **(E)** There is no effect of genotype on sleep duration upon silencing *IR56D*-expressing neurons (two-way ANOVA: F_1,200_= 0.5138, *P*=0.4743), but the effect of food treatment remains (two-way ANOVA: F_2,200_= 71.95, *P*<0.0001). Relative to agar, both genotypes significantly increase sleep duration when HxA is added to the diet (*IR56D*>imp*TNT*: *P*<0.0001; *IR56D*>*TNT*: *P*<0.0001), however it is significantly less than flies fed standard food (*IR56D*>imp*TNT*: *P*=0.0173; *IR56D*>*TNT*: *P*<0.0188). **(F)** There is no effect of genotype on the number of sleep episodes upon silencing *IR56D*-expressing neurons (two-way ANOVA: F_1,200_= 2.423, *P*=0.1211), but the effect of food treatment remains (two-way ANOVA: F_2,200_= 82.19, *P*<0.0001). Relative to agar, both genotypes significantly increase sleep bout number when HxA is added to the diet (*IR56D*>imp*TNT*: *P*<0.0001; *IR56D*>*TNT*: *P*<0.0001), however this is not different than when flies are fed standard food (*IR56D*>imp*TNT*: *P*=0.4342; *IR56D*>*TNT*: *P*=0.3257). **(G)** There is no effect of genotype on the length of each sleep episode upon silencing *IR56D*-expressing neurons (two-way ANOVA: F_1,200_= 2.407, *P*=0.1224), nor is there an effect of food treatment (two-way ANOVA: F_2,200_= 2.308, *P*=0.1021). **(H)** There is no effect of genotype on waking activity upon silencing *IR56D*-expressing neurons (two-way ANOVA: F_1,200_= 0.8303, *P*=0.3633), but the effect of food treatment remains (two-way ANOVA: F_2,200_= 19.77, *P*<0.0001). Relative to agar, there is no difference in waking activity upon the addition of HxA in control flies or after silencing IR56D-expressing neurons (*IR56D*>imp*TNT*: *P*=0.8707; *IR56D*>*TNT*: *P*=0.3591), but is significantly higher than on standard food (*IR56D*>imp*TNT*: *P*=0.0008; *IR56D*>*TNT*: *P*=0.0185). Bar graphs display mean ± s.e.m. Gray circles represent measurements of individual flies. A-D: N = 21-101; E-H: N = 32-39. ** *P*<0.01; *** *P*<0.001; **** *P*<0.0001.

## Discussion

Our findings reveal that dietary fatty acids promote sleep in flies fed a nutrient poor diet. Previous studies have found that starvation induces wakefulness and hyperactivity, suggesting flies forgo sleep in order to forage (Keene et al., 2010; G. Lee & Park, 2004; Mattaliano, Montana, Parisky, Littleton, & Griffith, 2007). This phenotype occurs across diverse *Drosophila melanogaster* laboratory strains and outbred lines (Brown et al., 2018; Masek et al., 2014), suggesting it is not an artifact of laboratory breeding. While previous work has shown that flies fed a diet of sugar alone sleep more than starved flies (Hasegawa et al., 2017; Keene et al., 2010; Sonn et al., 2018; B. Stahl, Slocumb, Chaitin, DiAngelo, & Keene, 2017), and that high calorie diets disrupt sleep (Catterson et al., 2010; Yamazaki et al., 2012), the contributions of different dietary macronutrients to sleep regulation has not been studied systematically. These findings suggest that fatty acids may provide an important component of the *Drosophila* diet that contributes to sleep regulation.

In diverse animal species, disruption of fatty acid metabolism has also been implicated in sleep regulation. For example, a mutation in an enzyme of the fatty acid β-oxidation pathway, *short-chain acylcoenzyme A dehydrogenase*, disrupts REM sleep in mice (Tafti et al., 2003). High fat diets can also disrupt fatty acid metabolism in flies (Heinrichsen et al., 2014), and have varied effects on sleep, resulting in an increase in sleep pressure, duration, and fragmentation (Grandner, Kripke, Naidoo, & Langer, 2010; Jenkins et al., 2006). In addition to sleep, high amounts of fat in the diet influence diverse traits, including triglyceride levels, longevity, as well as resistance to stress (Heinrichsen & Haddad, 2012). Overall, these findings contribute to a growing literature suggesting the type and quantity of individual macronutrients, including fats, have broad effects on sleep regulation and physiological adaptations (Grandner, Jackson, Gerstner, & Knutson, 2014; Lindseth, Lindseth, & Thompson, 2013).

Fatty acids, as well as esters and alcohols, are a by-product of yeast fermentation (Styger et al., 2011), and the presence of yeast is the primary attractant of *Drosophila* to fruit (Becher et al., 2012; Hoang, Kopp, & Chandler, 2015; Palanca, Gaskett, Günther, Newcomb, & Goddard, 2013). Hexanoic acid, in particular, is a by-product of human-made fermentation products, including wine and vinegar (Stökl et al., 2010). Fatty acids are also naturally produced by fruits consumed by *Drosophila*. For example, they are present in the banana during the later stages of ripening (Zhu et al., 2018). In fruit, fatty acids are metabolized to form esters (Balbontín et al., 2010; Qin et al., 2014), which are among the most behaviorally relevant compounds, as both the larval and adult olfactory systems are largely tuned to this class of volatiles (Grabe et al., 2018). Although much is known about the peripheral detection of these chemicals, it remains unknown how their consumption modulates behavior. Our study begins to elucidate this in regard to sleep regulation. Interestingly, the Morinda fruit, *M. citrofolia*, the primary food source of *Drosophila sechelia* is rich in fatty acids, primarily octanoic and hexanoic acids (Farine et al., 1996; Pino, Márquez, Quijano, & Castro, 2010). While this is reported to be toxic to several species of *Drosophila,* including *D. melanogaster* (Farine et al., 1996), our finding that HxA promotes sleep and that survivorship is enhanced in flies fed low doses of medium-chain fatty acids (Masek & Keene, 2013) raise the possibility that low doses of FAs promote survival.

Our findings revealed that dietary fatty acids ranging from 6-10 carbons promote sleep, yet have diverse effects on bout length, bout number, and waking activity, thereby raising the possibility that multiple cellular or neural mechanisms regulate fatty acid-dependent modulation of sleep duration. Dietary nutrients can be perceived via peripheral processing by the taste system as well as by taste-independent mechanisms (Lenard & Berthoud, 2008; Loper, La Sala, Dotson, & Steinle, 2015; Morton, Cummings, Baskin, Barsh, & Schwartz, 2006). In the taste system, thermogenetically activating sweet sensing neurons labeled by the *Gustatory receptor 64f* is sufficient to restore normal sleep to starved flies (Linford et al., 2015). However, whether flies lacking sugar receptors suppress sleep on an all sugar diet have been conflicting (Hasegawa et al., 2017; Keene et al., 2010; Linford et al., 2012). Our finding that flies with silenced fatty acid-sensing IR56D-expressing neurons, or a deletion of the IR56D gene, continue to increase sleep on a diet of HxA suggests the sleep-promoting effects of HxA are independent of the canonical taste pathway.

Given that sleep regulation by dietary fatty acids is independent of their role in taste regulation, it is possible that sleep regulation by dietary fatty acids occurs centrally. Multiple populations of central brain neurons that sense nutrient levels or modulate feeding behavior have been identified as essential for integrating sleep and activity with nutritional status (Chung et al., 2017; G. Lee & Park, 2004; Mattaliano, Montana, Parisky, Littleton, & Griffith, 2007; M. E. Yurgel et al., 2019). For example, the peptidergic *Leucokinin*-expressing neurons increase activity during starvation conditions, and when silenced, fail to suppress sleep (M. E. Yurgel et al., 2019). Further, overexpression of *short neuropeptide F*-expressing neurons suppress sleep and promote food consumption (Chen et al., 2013; K. S. Lee, You, Choo, Han, & Yu, 2004). Nutrient-sensitive central brain neurons, including the insulin producing cells and the sodium/glucose co-transporter SLC5A11 (Dus, Ai, & Suh, 2013; Nässel, Kubrak, Liu, Luo, & Lushchak, 2013), may also be involved in the regulation of sleep by dietary fatty acids. Further, *fatty acid binding proteins* (*Fabp)* that comprise a group of soluble protein transporters that bind small hydrophobic lipids and act as transporters have been implicated in sleep regulation raising the possibility that these are targets of dietary fatty acids(Gerstner et al 2011, 2017). It is also possible that the dietary regulation of sleep may occur via brain-periphery communication, as peripheral metabolic tissue, such as the fat body, has been previously shown to contribute to sleep regulation (Bennick et al., 2019; Thimgan, Suzuki, Seugnet, Gottschalk, & Shaw, 2010).

While consumption of dietary fatty acids increases survivorship in otherwise starved flies, induces a robust feeding response (Masek & Keene, 2013), and promotes sleep, the role of fatty acid consumption in fly behavior under natural conditions is poorly understood, thereby highlighting a critical need to understand the ethological relevance of laboratory rearing and behavioral analysis. Nevertheless, the robust changes to sleep duration and architecture upon consumption of diverse classes of fatty acids suggest they may play a critical role in regulating fly behavior. Taken together, these findings provide additional evidence that sleep in flies is regulated by caloric value and macronutrients within their diet and provide a platform for identifying internal fatty acid sensors.

## Materials and Methods

### *Drosophila* stocks and husbandry

Flies were grown and maintained on standard *Drosophila* food media (Bloomington Recipe, Genesee Scientific, San Diego, CA, USA). All experiments were performed in incubators (Powers Scientific, Warminster, PA, USA) on a 12:12 LD cycle at 25°C with a humidity of 55– 65%. The following fly strains were obtained from the Bloomington Stock Center: *w^1118^*(#5905); *IR56D*-GAL4 (#60708; (Koh et al., 2014)); UAS-imp*TNT* (#28840; (Sweeney et al., 1995)); and UAS-*TNT* (#28996; (Sweeney et al., 1995) wer obtained from the blooming stock center. For all experiments, mated females flies aged three-to-five days were used.

### Preparation of fatty acids

The following fatty acids were obtained from Sigma Aldrich (St Louis, MO, USA): propionic acid (3C; #402907), butyric acid (4C; #B103500), valeric acid (5C; #240370), hexanoic acid (6C; #21530), heptanoic acid (7C; #75190), octanoic acid (8C; #O3907), nonanoic acid (9C; #N5502), and decanoic acid (10C; #C1875). Unless otherwise indicated, all fatty acids were tested at a concentration of 0.4% and were dissolved in 1% agar (#BP1423, Fisher Scientific, Hampton, NH, USA). Once the 1% agar solution reached a boil, it was removed from heat and then fatty acid was added. This solution was stirred until the fatty acid was completely dissolved.

### Measurements of sleep and activity

Individual female flies aged 3-5 days were placed into clear plastic tubes containing standard food. Flies were acclimated to these experimental conditions for at least 24 hrs prior to the start of behavioral analysis. Measurements of sleep and waking activity were then measured over a 24 hr period starting at ZT0 using the *Drosophila* Locomotor Activity Monitor (DAM) System (Trikinetics, Waltham, MA, USA) as previously described (Hendricks et al., 2000; Shaw et al., 2000). At ZT0 the following day, flies were transferred to 0.4% HxA or agar, unless otherwise specified, and sleep and waking activity were again measured over a 24 hr period. For each individual fly, the DAM system measures activity by counting the number of infrared beam crossings over time. These activity data were then used to calculate sleep, defined as bouts of immobility of 5 min or more, using the *Drosophila* Sleep Counting Macro (Pfeiffenberger, Lear, Keegan, & Allada, 2010), from which sleep traits were then extracted.

### Statistical analysis

Measurements of sleep traits are presented as bar graphs showing mean ± standard error. A one-way or two-way analysis of variance (ANOVA) was used for comparisons between two or more treatments or two or more genotypes and two or more treatments, respectively. All post hoc analyses were performed using Tukey’s Honest Significant Difference test. A t-test was used for comparisons between two treatments. Statistical analyses and data presentation were performed using InStat software (GraphPad Software 8.0; San Diego, CA, USA).

## Acknowledgments

We would like to thank members of the Keene lab for technical assistance and helpful discussion. This work was funded by NIH grants R01 NS085252 and R01DC017390 to ACK, as well as support from FAU’s Jupiter Life Science Initiative.

## Supplementary Information

**Figure S1.**
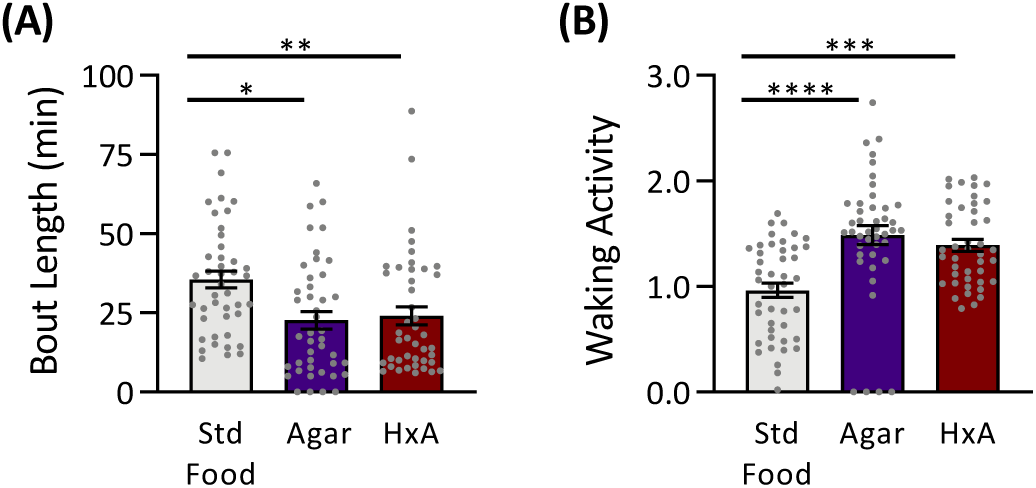
**Bout length and waking activity measurements from flies** that were transferred to their respective diet at ZT12. This data is a continuation from Figure 1, as sleep duration and bout number are shown in Figure 1H-I. **(A)** The length of each sleep episode is dependent on food treatment (one-way ANOVA: F_2,126_ = 6.438, P=0.0022). Flies fed standard food have significantly longer bout lengths than flies fed a diet of either agar (*P*=0.0039) or HxA (*P*=0.0118), which are not significantly different from each other (*P*=0.9296). **(B)** Waking activity is dependent on food treatment (one-way ANOVA: F_2,126_= 14.21, *P*<0.0001). Flies fed standard food are significantly less active than flies fed a diet of either agar (*P*<0.0001) or HxA (*P*=0.0002), which are not significantly different from each other (*P*=0.6349). Bar graphs display mean ± s.e.m. Gray circles represent measurements of individual flies.

**Figure S2.**
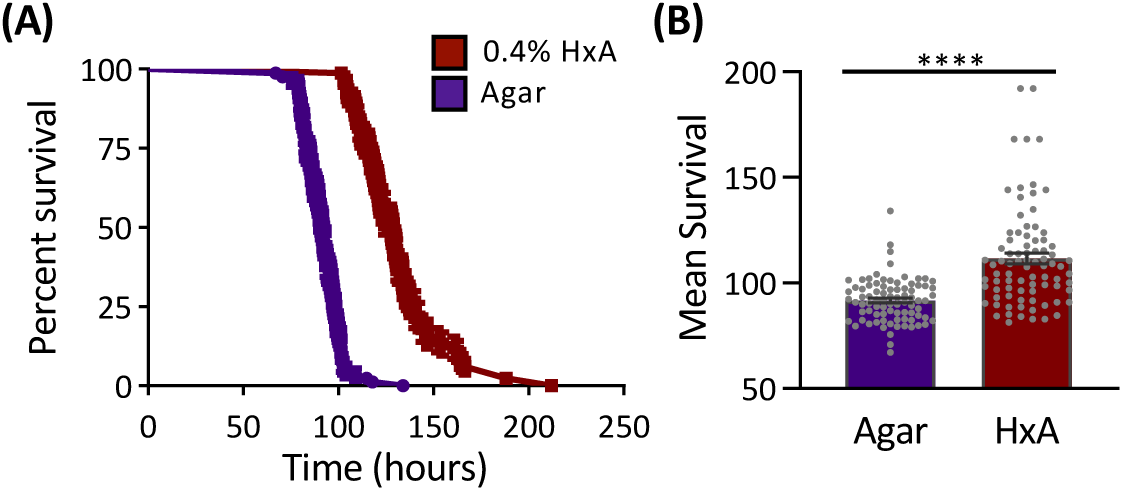
Dietary fatty acid promotes longevity. Flies fed 0.4% HxA survive significantly longer than flies fed agar alone (Log-Rank test: χ^2^= 58.58, d.f.=1, *P*<0.0001). **(A)** Survivorship curves showing the percentage of flies remaining alive as a function of the duration of starvation. **(B)** Mean survivorship of flies fed 0.4% HxA compared to agar alone. Survivorship was measured once flies were transferred to agar. N = 83. Bar graphs display mean ± s.e.m. Gray circles represent measurements of individual flies.

## References

Ahn, J.-E., Chen, Y., & Amrein, H. (2017). Molecular basis of fatty acid taste in *Drosophila*. ELife, 6, 1–21. https://doi.org/10.7554/elife.30115

Allada, R., & Siegel, J. M. (2008). Unearthing the phylogenetic roots of sleep. Current Biology, 18, 670–679. https://doi.org/10.1016/j.cub.2008.06.033

Alphen, B. van, Yap, M. H. W., Kirszenblat, L., Kottler, B., & Swinderen, B. van. (2013). A Dynamic Deep Sleep Stage in *Drosophila*. The Journal of Neuroscience, 33(16), 6917– 6927. https://doi.org/10.1523/JNEUROSCI.0061-13.2013

Arble, D. M., Bass, J., Behn, C. D., Butler, M. P., Challet, E., Czeisler, C., … Wright, K. P. (2015). Impact of Sleep and Circadian Disruption on Energy Balance and Diabetes: A Summary of Workshop Discussions. Sleep, 38(12), 1849–1860. https://doi.org/10.5665/sleep.5226

Balbontín, C., Gaete-Eastman, C., Fuentes, L., Figueroa, C. R., Herrera, R., Manriquez, D., … Moya-León, M. A. (2010). *VpAAT1*, a gene encoding an alcohol acyltransferase, is involved in ester biosynthesis during ripening of mountain papaya fruit. Journal of Agricultural and Food Chemistry, 58(8), 5114–5121. https://doi.org/10.1021/jf904296c

Becher, P. G., Flick, G., Rozpedowska, E., Schmidt, A., Hagman, A., Lebreton, S., … Bengtsson, M. (2012). Yeast, not fruit volatiles mediate *Drosophila melanogaster* attraction, oviposition and development. Functional Ecology, 26(4), 822–828. https://doi.org/10.1111/j.1365-2435.2012.02006.x

Bennick, R. A., Shah, K. D., Burns, C., Yurgel, M. E., Keene, A. C., Brown, E. B., & DiAngelo, J. R. (2019). *Ade2* functions in the *Drosophila* fat body To promote sleep. G3, 8(11), 3385– 3395. https://doi.org/10.1534/g3.118.200554

Brown, E. B., Torres, J., Bennick, R. A., Rozzo, V., Kerbs, A., Diangelo, J. R., & Keene, A. C. (2018). Variation in sleep and metabolic function is associated with latitude and average temperature in *Drosophila melanogaster*. Ecology and Evolution, 8, 4084– 4097. https://doi.org/10.1002/ece3.3963

Capellini, I., Barton, R. A., McNamara, P., Preston, B. T., & Nunn, C. L. (2008). Phylogenetic analysis of the ecology and evolution of mammalian sleep. Evolution, 62, 1764–1776. https://doi.org/10.1111/j.1558-5646.2008.00392.x

Catterson, J. H., Knowles-Barley, S., James, K., Heck, M. M. S., Harmar, A. J., & Hartley, P. S. (2010). Dietary modulation of *Drosophila* sleep-wake behaviour. PLoS ONE, 5(8). https://doi.org/10.1371/journal.pone.0012062

Chakravarti, L., Moscato, E. H., & Kayser, M. S. (2017). Unraveling the neurobiology of sleep and sleep disorders using Drosophila. 121, 253–285. https://doi.org/10.1016/bs.ctdb.2016.07.010

Chen, W., Shi, W., Li, L., Zheng, Z., Li, T., Bai, W., & Zhao, Z. (2013). Regulation of sleep by the short neuropeptide F (sNPF) in *Drosophila melanogaster*. Insect Biochemistry and Molecular Biology, 43(9), 809–819. https://doi.org/10.1016/j.ibmb.2013.06.003

Chung, B. Y., Ro, J., Hutter, S. A., Miller, K. M., Guduguntla, L. S., Kondo, S., & Pletcher, S. D. (2017). *Drosophila* Neuropeptide F signaling independently regulates feeding and sleep-wake behavior. Cell Reports, 19(12), 2441–2450. https://doi.org/10.1016/j.celrep.2017.05.085

Donlea, J. M. (2017). Neuronal and molecular mechanisms of sleep homeostasis. Current Opinion in Insect Science, 24, 51–57. https://doi.org/10.1016/j.cois.2017.09.008

Dus, M., Ai, M., & Suh, G. S. B. (2013). Taste-independent nutrient selection is mediated by a brain-specific Na +/solute co-transporter in *Drosophila*. Nature Neuroscience, 16(5), 526– 528. https://doi.org/10.1038/nn.3372

Farine, J., Legal, L. U. C., Moreteauf, B., I, J. L. E. Q., Laboratoire, I., Recherches, D., … F-, G. (1996). Volatile components of ripe fruits of *Morinda citrifolia* and their effects on *Drosophila*. Science, 4(2), 433–438. https://doi.org/10.1016/0031-9422(95)00455-6

Faville, R., Kottler, B., Goodhill, G., Shaw, P. J., & van Swinderen, B. (2015). How deeply does your mutant sleep? Probing arousal to better understand sleep defects in Drosophila. Science Reports, 13(5), 8454. https://doi.org/10.1038/srep08454

Fischler, W., Kong, P., Marella, S., & Scott, K. (2007). The detection of carbonation by the *Drosophila* gustatory system. Nature, 448(7157), 1054–1057. https://doi.org/10.1038/nature06101

Grabe, V., Weißflog, J., Svatoš, A., Dweck, H. K. M., Knaden, M., Retzke, T., … Hansson, B. S. (2018). The olfactory logic behind fruit odor preferences in larval and adult *Drosophila*. Cell Reports, 23(8), 2524–2531. https://doi.org/10.1016/j.celrep.2018.04.085

Grandner, M. A., Jackson, N., Gerstner, J. R., & Knutson, K. L. (2014). Sleep symptoms associated with intake of specific dietary nutrients. Journal of Sleep Research, 23(1), 22– 34. https://doi.org/10.1111/jsr.12084

Grandner, M. A., Kripke, D. F., Naidoo, N., & Langer, R. D. (2010). Relationships among dietary nutrients and subjective sleep, objective sleep, and napping in women. Sleep Medicine, 11(2), 180–184. https://doi.org/10.1016/j.sleep.2009.07.014

Griffith, L. C. (2013). Neuromodulatory control of sleep in *Drosophila melanogaster*: Integration of competing and complementary behaviors. Current Opinion in Neurobiology, 23(5), 819– 823. https://doi.org/10.1016/j.conb.2013.05.003

Hasegawa, T., Tomita, J., Hashimoto, R., Ueno, T., Kume, S., & Kume, K. (2017). Sweetness induces sleep through gustatory signalling independent of nutritional value in a starved fruit fly. Scientific Reports, 7(1), 1–9. https://doi.org/10.1038/s41598-017-14608-1

Heinrichsen, E. T., & Haddad, G. G. (2012). Role of high-fat diet in stress response of *Drosophila*. PLoS ONE, 7(8), 3–10. https://doi.org/10.1371/journal.pone.0042587

Heinrichsen, E. T., Zhang, H., Robinson, J. E., Ngo, J., Diop, S., Bodmer, R., … Haddad, G. G. (2014). Metabolic and transcriptional response to a high-fat diet in *Drosophila melanogaster*. Molecular Metabolism, 3(1), 42–54. https://doi.org/10.1016/j.molmet.2013.10.003

Hendricks, J. C., Finn, S. M., Panckeri, K. A., Chavkin, J., Williams, J. A., Sehgal, A., & Pack, A. I. (2000). Rest in *Drosophila* is a sleep-like state. Neuron, 25, 129–138. https://doi.org/10.1016/S0896-6273(00)80877-6

Hoang, D., Kopp, A., & Chandler, J. A. (2015). Interactions between *Drosophila* and its natural yeast symbionts - Is *Saccharomyces cerevisiae* a good model for studying the fly-yeast relationship? PeerJ, 3, e1116. https://doi.org/10.7717/peerj.1116

Jenkins, J. B., Omori, T., Guan, Z., Vgontzas, A. N., Bixler, E. O., & Fang, J. (2006). Sleep is increased in mice with obesity induced by high-fat food. Physiology and Behavior, 87(2), 255–262. https://doi.org/10.1016/j.physbeh.2005.10.010

Joiner, W. J. (2016). Unraveling the evolutionary determinants of sleep. Current Biology, 26(20), 1073–1087. https://doi.org/10.1016/j.cub.2016.08.068

Keene, A. C., & Duboue, E. R. (2018). The origins and evolution of sleep. The Journal of Experimental Biology, 12(221), jeb159533. https://doi.org/10.1242/jeb.159533

Keene, A. C., Duboué, E. R., McDonald, D. M., Dus, M., Suh, G. S. B., Waddell, S., & Blau, J. (2010). Clock and cycle limit starvation-induced sleep loss in *Drosophila*. Current Biology, 20(13), 1209–1215. https://doi.org/10.1016/j.cub.2010.05.029

Koh, T. W., He, Z., Gorur-Shandilya, S., Menuz, K., Larter, N. K., Stewart, S., & Carlson, J. R. (2014). The *Drosophila* IR20a clade of ionotropic receptors are candidate taste and pheromone receptors. Neuron, 83(4), 850–865. https://doi.org/10.1016/j.neuron.2014.07.012

Lee, G., & Park, J. H. (2004). Hemolymph sugar homeostasis and starvation-induced hyperactivity affected by genetic manipulations of the adipokinetic hormone-encoding gene in *Drosophila melanogaster*. Genetics, 167, 311–323. https://doi.org/10.1534/genetics.167.1.311

Lee, K. S., You, K. H., Choo, J. K., Han, Y. M., & Yu, K. (2004). *Drosophila* short neuropeptide F regulates food intake and body size. Journal of Biological Chemistry, 279(49), 50781–50789. https://doi.org/10.1074/jbc.M407842200

Lenard, N. R., & Berthoud, H.-R. (2008). Central and peripheral regulation of food intake and physical activity: Pathways and Genes. Obesity, 16(S3), S11–S22. https://doi.org/10.1038/oby.2008.511

Lindseth, G., Lindseth, P., & Thompson, M. (2013). Nutritional effects on sleep. Western Journal of Nursing Research, 35(4), 497–513. https://doi.org/10.1177/0193945911416379

Linford, N. J., Chan, T. P., & Pletcher, S. D. (2012). Re-patterning sleep architecture in *Drosophila* through gustatory perception and nutritional quality. PLoS Genetics, 8(5), e1002668. https://doi.org/10.1371/journal.pgen.1002668

Linford, N. J., Ro, J., Chung, B. Y., & Pletcher, S. D. (2015). Gustatory and metabolic perception of nutrient stress in *Drosophila*. Proceedings of the National Academy of Sciences of the United States of America, 112(8), 2587–2592. https://doi.org/10.1073/pnas.1401501112

Loper, H. B., La Sala, M., Dotson, C., & Steinle, N. (2015). Taste perception, associated hormonal modulation, and nutrient intake. Nutrition Reviews, 73(2), 83–91. https://doi.org/10.1093/nutrit/nuu009

Masek, P., & Keene, A. C. (2013). *Drosophila* fatty acid taste signals through the PLC pathway in sugar-sensing neurons. PLoS Genetics, 9(9), e1003710. https://doi.org/10.1371/journal.pgen.1003710

Masek, P., Reynolds, L. a, Bollinger, W. L., Moody, C., Mehta, A., Murakami, K., … Keene, A. C. (2014). Altered regulation of sleep and feeding contribute to starvation resistance in *Drosophila*. The Journal of Experimental Biology, 217(17), 3122–3132. https://doi.org/10.1242/jeb.103309

Mattaliano, M. D., Montana, E. S., Parisky, K. M., Littleton, J. T., & Griffith, L. C. (2007). The *Drosophila* ARC homolog regulates behavioral responses to starvation. Molecular and Cellular Neuroscience, 36(2), 211–21. https://doi.org/10.1016/j.mcn.2007.06.008

Morton, G. J., Cummings, D. E., Baskin, D. G., Barsh, G. S., & Schwartz, M. W. (2006). Central nervous system control of food intake and body weight. Nature, 443(7109), 289–295. https://doi.org/10.1038/nature05026

Nässel, D. R., Kubrak, O. I., Liu, Y., Luo, J., & Lushchak, O. V. (2013). Factors that regulate insulin producing cells and their output in *Drosophila*. Frontiers in Physiology, 4(252), 1–12. https://doi.org/10.3389/fphys.2013.00252

Palanca, L., Gaskett, A. C., Günther, C. S., Newcomb, R. D., & Goddard, M. R. (2013). Quantifying variation in the ability of yeasts to attract *Drosophila melanogaster*. PLoS ONE, 8(9), 1–10. https://doi.org/10.1371/journal.pone.0075332

Pfeiffenberger, C., Lear, B. C., Keegan, K. P., & Allada, R. (2010). Locomotor activity level monitoring using the *Drosophila* activity monitoring (DAM) system. Cold Spring Harbor Protocols, 5. https://doi.org/10.1101/pdb.prot5518

Pino, J. A., Márquez, E., Quijano, C. E., & Castro, D. (2010). Volatile compounds in noni (Morinda citrifolia L.) at two ripening stages. Ciência e Tecnologia de Alimentos, 30(1), 183–187. https://doi.org/10.1590/s0101-20612010000100028

Qin, G., Tao, S., Zhang, H., Huang, W., Wu, J., Xu, Y., & Zhang, S. (2014). Evolution of the aroma volatiles of pear fruits supplemented with fatty acid metabolic precursors. Molecules, 19(12), 20183–20196. https://doi.org/10.3390/molecules191220183

Sehgal, A., & Mignot, E. (2011). Genetics of sleep and sleep disorders. Cell, 146(2), 194–207. https://doi.org/10.1016/j.cell.2011.07.004

Shaw, P. J., Cirelli, C., Greenspan, R. J., & Tononi, G. (2000). Correlates of sleep and waking in *Drosophila melanogaster*. Science, 287, 1834–1837. https://doi.org/10.1126/science.287.5459.1834

Siegel, J. M. (2005). Clues to the functions of mammalian sleep. Nature, 437, 1264–1271. https://doi.org/10.1038/nature04285

Sonn, J. Y., Lee, J., Sung, M. K., Ri, H., Choi, J. K., Lim, C., & Choe, J. (2018). Serine metabolism in the brain regulates starvation-induced sleep suppression in *Drosophila melanogaster*. Proceedings of the National Academy of Sciences of the United States of America, 201719033. https://doi.org/10.1073/pnas.1719033115

Stahl, B., Slocumb, M., Chaitin, H., DiAngelo, J., & Keene, A. (2017). Sleep-dependent modulation of metabolic rate In *Drosophila*. Sleep, 40(8), zsx084. https://doi.org/10.1093/sleep/zsx084

Stökl, J., Strutz, A., Dafni, A., Svatos, A., Doubsky, J., Knaden, M., … Stensmyr, M. C. (2010). A deceptive pollination system targeting Drosophilids through olfactory mimicry of yeast. Current Biology, 20(20), 1846–1852. https://doi.org/10.1016/j.cub.2010.09.033

Styger, G., Jacobson, D., & Bauer, F. F. (2011). Identifying genes that impact on aroma profiles produced by *Saccharomyces cerevisiae* and the production of higher alcohols. Applied Microbiology and Biotechnology, 91(3), 713–730. https://doi.org/10.1007/s00253-011-3237-z

Sweeney, S. T., Broadie, K., Keane, J., Niemann, H., & Kane, C. J. O. (1995). Targeted expression of tetanus toxin light chain in *Drosophila* specifically eliminates synaptic transmission and causes behavioral defects. Neuron, 14, 341–351. https://doi.org/10.1016/0896-6273(95)90290-2

Tafti, M., Petit, B., Chollet, D., Neidhart, E., De Bilbao, F., Kiss, J. Z., … Franken, P. (2003). Deficiency in short-chain fatty acid β-oxidation affects theta oscillations during sleep. Nature Genetics, 34(3), 320–325. https://doi.org/10.1038/ng1174

Tan, X., Alén, M., Cheng, S. M., Mikkola, T. M., Tenhunen, J., Lyytikäinen, A., … Cheng, S. (2015). Associations of disordered sleep with body fat distribution, physical activity and diet among overweight middle-aged men. Journal of Sleep Research, 24(4), 414–424. https://doi.org/10.1111/jsr.12283

Tauber, J., Brown, E., Li, Y., Yurgel, M., Masek, P., & Keene, A. (2017). A subset of sweet-sensing neurons identified by IR56d are necessary and sufficient for fatty acid taste. PLoS Genetics, 13(11), e1007059. https://doi.org/https://doi.org/10.1101/174623

Tauber, J. M., Brown, E. B., Li, Y., Yurgel, M. E., Masek, P., & Keene, A. C. (2017). A subset of sweet-sensing neurons identified by *IR56d* are necessary and sufficient for fatty acid taste. PLOS Genetics, 13(11), e1007059. https://doi.org/10.1371/journal.pgen.1007059

Thimgan, M. S., Suzuki, Y., Seugnet, L., Gottschalk, L., & Shaw, P. J. (2010). The Perilipin homologue, *Lipid Storage Droplet 2*, regulates sleep homeostasis and prevents learning impairments following sleep los*s*. PLOS Biology, 8(8), e1000466. https://doi.org/10.1371/journal.pbio.1000466

Toshima, N., & Tanimura, T. (2012). Taste preference for amino acids is dependent on internal nutritional state in *Drosophila melanogaster*. Journal of Experimental Biology, 215(16), 2827–2832. https://doi.org/10.1242/jeb.069146

Yamazaki, M., Tomita, J., Takahama, K., Ueno, T., Mitsuyoshi, M., Sakamoto, E., … Kume, K. (2012). High calorie diet augments age-associats sleep impairment in *Drosophila*. Biochemical and Biophysical Research Communications, 417(2), 812–816. https://doi.org/10.1016/j.bbrc.2011.12.041

Yurgel, M. E., Kakad, P., Zandawala, M., Nässel, D. R., Godenschwege, T. A., & Keene, A. C. (2019). A single pair of leucokinin neurons are modulated by feeding state and regulate sleep–metabolism interactions. PLoS Biology, 17(2), e2006409. https://doi.org/10.1371/journal.pbio.2006409

Yurgel, M., Masek, P., DiAngelo, J. R., & Keene, A. (2014). Genetic dissection of sleep-metabolism interactions in the fruit fly. Journal of Comparative Physiology A: Neuroethology,, Sensory, Neural, and Behavioral Physiology, 201(9), 869–77. https://doi.org/10.1007/s00359-014-0936-9

Zappia, G., Neagu-Maier, G. L., Sprecher, S. G., Sahai, S. Y., Silbering, A. F., Münch, D., … Croset, V. (2018). An expression atlas of variant ionotropic glutamate receptors identifies a molecular basis of carbonation sensing. Nature Communications, 9(1), 4252. https://doi.org/10.1038/s41467-018-06453-1

Zhang, Y. V, Ni, J., & Montell, C. (2013). The molecular basis for attractive salt-taste coding in *Drosophila*. Science (New York, N.Y.), 340(6138), 1334–1338. https://doi.org/10.1126/science.1234133

Zhu, X., Li, Q., Li, J., Luo, J., Chen, W., & Li, X. (2018). Comparative study of volatile compounds in the fruit of two banana cultivars at different ripening stages. Molecules, 23(10), 2456. https://doi.org/10.3390/molecules23102456

